# High-Intensity Interval Training Attenuates Impairment in Regulatory Protein Machinery of Mitochondrial Quality Control in Skeletal Muscle of Diet-Induced Obese Mice

**DOI:** 10.1101/2023.06.28.546902

**Authors:** James B. Tincknell, Benjamin Kugler, Haley Spicuzza, Huimin Yan, Tongjian You, Kai Zou

## Abstract

Mitochondrial quality control processes are essential in governing mitochondrial integrity and function. The purpose of the study was to examine the effects of 10 weeks of HIIT on the regulatory protein machinery of skeletal muscle mitochondrial quality control and whole-body glucose homeostasis in diet-induced obese mice. Male C57BL/6 mice were randomly assigned to a low-fat diet (LFD) or high-fat diet (HFD) group. After 10 weeks, HFD-fed mice were divided into sedentary and HIIT (HFD+HIIT) groups and remained on HFD for another 10 weeks (n=9/group). Graded exercise test, glucose and insulin tolerance tests, mitochondrial respiration and regulatory protein markers of mitochondrial quality control processes were determined by immunoblots. Ten weeks of HIIT enhanced ADP-stimulated mitochondrial respiration in diet-induced obese mice (P < 0.05) but did not improve whole-body insulin sensitivity. Importantly, the ratio of Drp1(Ser^616^) over Drp1(Ser^637^) phosphorylation, an indicator of mitochondrial fission, was attenuated in HFD-HIIT compared to HFD (−35.7%, P < 0.05). Regarding autophagy, skeletal muscle p62 content was lower in HFD group than LFD group (−35.1%, P < 0.05), however, such reduction was disappeared in HFD+HIIT group. In addition, LC3B II/I ratio was higher in HFD than LFD group (15.5%, P < 0.05) but was ameliorated in HFD+HIIT group (−29.9%, P < 0.05). Overall, our study demonstrated that 10 weeks of HIIT was effective in improving skeletal muscle mitochondrial respiration and the regulatory protein machinery of mitochondrial quality control in diet-induced obese mice through the alterations of mitochondrial fission protein Drp1 activity and p62/LC3B-mediated regulatory machinery of autophagy.

## INTRODUCTION

The prevalence of obesity has continued to rise exponentially in the U.S., placing a significant burden on the healthcare system (Ward et al. 2019). Obesity is accompanied with a myriad of metabolic perturbations that contribute to an increased risk for chronic diseases, including type 2 diabetes, cardiovascular diseases, and cancer (Grundy 2000). Skeletal muscle is responsible for approximately 70-80% of whole-body glucose disposal under insulin-stimulated conditions (DeFronzo and Tripathy 2009). Therefore, dysregulation of skeletal muscle metabolism can strongly influence whole-body glucose homeostasis and insulin sensitivity.

Mitochondria play an integral role in regulating substrate metabolism and maintaining cellular homeostasis in skeletal muscle (Hood et al. 2019). These organelles form interconnected networks, which are dynamically regulated by a series of cellular processes (e.g., mitochondrial biogenesis, mitochondrial dynamics (fission and fusion), and mitophagy/autophagy), known as mitochondrial quality control (Slavin et al. 2022). In brief, mitochondrial fusion is the process that two mitochondria join at the outer and inner membrane, while mitochondrial fission is the process when mitochondrial networks are segregated into separated mitochondria (Twig and Shirihai 2011). Lastly, mitophagy selectively removes fragmented and damaged mitochondria (Twig and Shirihai 2011). The orchestrated balance between these processes is crucial for maintaining mitochondrial fitness and overall skeletal muscle health (Slavin et al. 2022).

Emerging studies (Axelrod et al. 2021; Greene et al. 2015; Leduc-Gaudet et al. 2018; Li et al. 2018; Liu et al. 2014), including ours (Gundersen et al. 2020; Kugler et al. 2020), have reported that skeletal muscle mitochondrial quality control was imbalanced in a manner that shifts towards excessive mitochondrial fission with impaired mitophagy/autophagy flux, resulting in the accumulation of damaged mitochondria and compromised mitochondrial fitness in skeletal muscle from both humans with insulin resistance and diet-induced insulin-resistant rodents. Altogether, the impairment in the cellular processes of mitochondrial quality control has been linked to the development of insulin resistance and type 2 diabetes (Fealy et al. 2018).

Chronic exercise training has been recommended as an effective strategy to improve skeletal muscle insulin sensitivity and whole-body metabolic health in obesity (Di Meo et al. 2017; Hawley and Lessard 2008; Richter and Hargreaves 2013; Venables and Jeukendrup 2008). Among various exercise modalities, high-intensity interval training (HIIT) has emerged as one of the most popular choices due to its time-efficient nature that fits the “on-the-go” mentality of modern society, as well as its comparable cardiometabolic benefits to those resulting from moderate-intensity continuous training (MICT) (Burgomaster et al. 2008; MacInnis and Gibala 2017; Poon et al. 2022; Su et al. 2019). While the effectiveness of HIIT on whole-body cardiometabolic health and skeletal muscle mitochondrial function in obesity has been well established (Batacan et al. 2017; Batterson et al. 2023; Campbell et al. 2019; Chavanelle et al. 2017), our knowledge with regards to how mitochondrial quality control processes may be regulated by HIIT is very limited. It seems the emphasis of recent research studies in the adaptations of mitochondrial quality control to exercise training has been almost exclusively on MICT (Axelrod et al. 2019; Fealy et al. 2014; Heo et al. 2021; Kugler et al. 2021b). Regarding HIIT, individual processes of mitochondrial quality control has been explored separately in several studies using short-term HIIT protocols (e.g., 2 weeks) (Batterson et al. 2023; MacInnis et al. 2017). Given that different exercise intensities and durations can lead to diverse mitochondrial adaptations in skeletal muscle (Dudley et al. 1982), it is important to examine the complete profile of mitochondrial quality control processes in adaptations to HIIT and how these adaptations are associated with cardiometabolic improvements. Therefore, the purpose of this study was to examine the effects of 10 weeks of HIIT intervention on the regulatory machinery of skeletal muscle mitochondrial quality control and whole-body glucose homeostasis in diet-induced obese mice. We hypothesized that high-fat diet feeding would impair mitochondrial respiration and the regulatory machinery of mitochondrial quality control towards pro-fission state, and induce whole-body insulin resistance, whereas 10 weeks of HIIT would attenuate the imbalance of mitochondrial dynamics and rescue mitochondrial dysfunction and whole-body insulin resistance in diet-induced obese mice.

## MATERIALS AND METHODS

### Animal Care

All experiments and procedures for animal use were approved by the Institutional Animal Care and Use Committee (IACUC) of the University of Massachusetts Boston (Protocol number: 2018193). Male C57/BL-6 mice were purchased (Jackson Laboratories, Bar Harbor, ME) at 8-weeks of age and acclimatized to the animal facility for 1-week before any experiment. Animals were housed in a temperature-controlled (22°C) environment and maintained on a 12:12 hour light-dark cycle with food and water provided ad libitum. Starting at 9 weeks of age, mice were randomly divided into either a high-fat diet (HFD, 45% kcal by fat, D12451, Research Diets, New Brunswick, NJ) or a low-fat diet group (LFD, 10% kcal by fat, D12450J, Research Diets, New Brunswick, NJ). After 10 weeks of diet intervention, mice in the HFD group were further randomly divided into either sedentary or HIIT groups for an additional 10 weeks. Therefore, the study consists of 3 groups: LFD, HFD, and HFD+HIIT (n = 9/group). Body weight and food consumption were weighed and recorded every week.

### Exercise Exhaustion Test

The protocol was based on previous works (Martinez-Huenchullan et al. 2018; Martinez-Huenchullan et al. 2019). Briefly, all mice were acclimatized to the Columbus Exer 3/6 Treadmill (Columbus, OH) for 3 days before the test. To assess the efficacy of the training protocol, an exercise performance test was performed before training commences to establish baseline measures and at the end of training (10 weeks).

Mice began treadmill running at a speed of 8 m/min. Speed was then increased at a rate of 1 m/min per 2 minutes of exercise until fatigued. Fatigue was defined as the inability of the animal to reach the end of the lane after encouragement with five mechanical stimuli (soft brush) delivered within one minute. Electric shock stimulation was not utilized during exercise capacity testing to avoid muscle damage. When fatigue was accomplished, speed was recorded as the maximal running speed (100%). The exercise prescription was the same for mice with less than 5% variability of maximal running speed.

### HIIT Training

The HIIT training protocol was adapted from previous works (Martinez-Huenchullan et al. 2018; Martinez-Huenchullan et al. 2019) with the following modifications. Mice participated in a warm-up for 5 consecutive minutes at 5 m/min. After the warm-up period, mice exercised at ∼85-90% maximal speed (as determined at baseline). There was a total of 6 bouts broken down into 3 minutes of exercise followed by a 2-minute active recovery at ∼50% maximal speed – totaling 30 minutes. After completing 6 bouts, there was a 5-minute cool-down period with a speed set at 5 m/min. Mice assigned to the HFD+HIIT group were exposed to a total of 10 weeks of training consisting of treadmill running for 3 non-consecutive days per week. The cages housing sedentary control group were placed near the treadmill without food and water during each training session.

### Glucose and Insulin Tolerance Tests

Glucose and insulin tolerance tests (GTT and ITT) were measured as previously described (Kugler et al. 2021a) during the last week of exercise training with 5 days apart to minimize the interference between the two tests. Prior to GTT and ITT, mice were fasted for 6 hours and given an intraperitoneal injection of either glucose (1 mg/kg body weight) or insulin (1 U/kg body weight). Blood samples were collected from the tail vein and assessed for glucose concentration before injection (0 min) and 30, 60, and 90 minutes (for ITT)/120 minutes (for GTT) post-injection using a Contour next blood glucometer (Ascensia Diabetes Care, Parsippany, NJ). The area under the curve (AUC) and inverse AUC were calculated using the trapezoid rule during the GTT and ITT, respectively (Nagy and Einwallner 2018).

### Tissue Collection

Forty-eight hours after the final bout of HIIT, mice were euthanized using CO_2_ asphyxiation/cervical dislocation. Blood was collected immediately via vena cava puncture and centrifuged for 15 minutes at 2,500 rpm at 4°C to collect serum. Freshly dissected quadriceps were collected and placed in ice-cold mitochondrial isolation buffer for mitochondrial respiration assay (Frisard et al. 2015). Gastrocnemius muscles were collected, snap-frozen in liquid nitrogen, and stored at −80°C for subsequent immunoblot analysis.

### Mitochondrial Respiration

Seahorse XFp (Agilent Technologies, Santa Clara, CA) Analyzer was used to determine mitochondria respiratory rates by measuring oxygen consumption rates (OCR). Mitochondria were isolated from freshly dissected quadriceps muscle, as previously described (Frisard et al. 2015). After mitochondria isolation, protein concentration was determined using a Pierce BCA protein assay kit (Thermo Fisher Scientific, Waltham, MA). Isolated mitochondria (3.5 μg/well) were then used in the presence of 10mM pyruvate and 5mM malate to determine mitochondrial respiration rates using a Seahorse XFp Extracellular Flux Analyzer (Agilent Technologies, Santa Clara, CA), as previously described (Kugler et al. 2021a). ADP (5mM), oligomycin (2μM), carbonyl cyanide-4 phenylhydrazone (FCCP, 4 μM), and antimycin (4 μM) were successively injected to measure OCR for different respiratory states. All data were analyzed using the Agilent Seahorse Wave software.

### Immunoblot Analyses

Gastrocnemius muscles were mechanically homogenized using a handheld homogenizer (Omni International, Kennesaw, Georgia) in ice-cold buffer (0.1% SDS, 0.1% sodium deoxycholate, 1% Triton X, 50mM Tris-HCl, 5mM EDTA, and 150mM NaCl,) supplemented with protease and phosphatase inhibitors (Thermo Scientific Inc., Waltham, MA). Protein concentrations were determined by Pierce BCA Protein Assay Kit (Thermo Fisher Scientific, Waltham, MA), and equal amounts of protein were subjected to SDS-PAGE using 4-20% gradient polyacrylamide gels (Bio-Rad, Hercules, CA) and transferred to a nitrocellulose membrane. Membranes were probed overnight with antibodies recognizing Voltage-dependent anion channel (VDAC, cat# 4661S), phosphorylated Drp1 Ser^616^ (cat# 3455S), phosphorylated Drp1 Ser^637^(cat# 6319S), Drp1 (cat# 8570S), Opa1 (cat# 67589S), LC3B (cat# 2775S), β-Actin (#3700S), GAPDH (cat# 2118S) (Cell Signaling, Danvers, MA), Fis1 (cat # sc-376447), MFF (cat# sc-398617) (Santa Cruz Biotechnology, Dallas, TX), OXPHOS Cocktail (cat# ab110413), Citrate Synthase (cat# ab96600), Mfn1 (cat# ab104274), Mfn2 (cat# ab56889), P62 (cat# ab56416), Parkin (cat# ab77924), PGC1-α (cat# ab106814) (Abcam, Cambridge, MA). Membranes were probed with an IRDye secondary antibody (Li-Cor, Lincoln, NE) and quantified using Odyssey CLx software (Li-Cor, Lincoln, NE). Data were normalized to β-Actin or GAPDH (for OXPHOS blot) protein expression.

### Statistical Analyses

All data was expressed as Mean ± SEM. Group-by-time interaction for GTT and ITT were analyzed using One-way repeated measures of analysis of variance (ANOVA). One-way ANOVA was used to assess any between-group differences. If a statistically significant main effect is determined, Tukey post hoc tests were used in order to establish group differences. Paired t-tests were used to assess pre-post differences in maximal speed. Pearson correlations were used to determine if there is a correlation between the expression of mitochondrial protein markers and measures of whole-body glucose metabolism. Based on the blood glucose result from a previously published study (Martinez-Huenchullan et al. 2019), in order to yield >80% power to detect the effect size [(7.7-7.0)/0.5=1.4] with 95% confidence, we will use samples of size n=9/group. Statistical significance was set at P < 0.05. IBM® SPSS Statistics 25.0 was used to analyze the data.

## RESULTS

### Effects of HIIT on body weight, whole-body insulin sensitivity, and exercise capacity in HFD-induced obese mice

We first verified that HFD resulted in obesity and insulin resistance. Twenty weeks of HFD resulted in significantly higher body weight and inguinal fat pad weight (P < 0.05, Table 1) than LFD. Importantly, HFD-fed mice that received 10 weeks of HIIT during HFD feeding displayed blunted body weight and fat pad weight gains when compared to the HFD group (−14.2% and −22.9%, P < 0.05, Table 1), while remained significantly higher than the LFD group (P < 0.05, Table 1). Mice fed with HFD had higher food consumption (g/day) than mice fed with LFD (P < 0.05, Table 1). No difference in absolute skeletal muscle or heart weight among groups. However, when normalized to body weight, gastrocnemius, TA, and heart weights were significantly lower in HFD-fed mice compared to LFD-fed mice, regardless of exercise training status (P < 0.05, Table 1).

**Table 1.** Animal phenotypic characteristics

|  | LFD | HFD | HFD+HIIT |
| --- | --- | --- | --- |
| Pre-HIIT Body weight (g) | 28.0±0.5 | 33.1±1.3* | 35.8±1.5* |
| Post-HIIT Body weight (g) | 30.9±0.7 | 46.5±1.5* | 39.4±1.6*<sup>#</sup> |
| Food consumption (g/day) | 2.97±0.05 | 3.14±0.12* | 3.09±0.06* |
| Tissue weight (g) |  |  |  |
| Gastrocnemius | 0.168±0.006 | 0.172±0.008 | 0.173±0.007 |
| Tibialis Anterior | 0.082±0.002 | 0.075±0.008 | 0.078±0.010 |
| Heart | 0.159±0.005 | 0.158±0.005 | 0.160±0.008 |
| Inguinal Fat Pad | 0.438±0.086 | 2.641±0.155* | 2.037±0.214*<sup>#</sup> |
| Tissue weight (mg/g of BW) |  |  |  |
| Gastrocnemius | 5.43±0.14 | 3.73±0.21*<sup>#</sup> | 4.32±0.24* |
| Tibialis Anterior | 2.67±0.09 | 1.61±0.17* | 1.95±0.26* |
| Heart | 5.15±0.17 | 3.42±0.16* | 4.03±0.29* |
| Inguinal Fat Pad | 15.27±2.98 | 57.14±5.41* | 51.72±4.58*<sup>#</sup> |

Data are presented as Mean ± SEM. BW: Body Weight \* P<0.05 vs. LFD, <sup>#</sup> P<0.05 vs. HFD.

To assess whole-body glucose homeostasis and insulin sensitivity, metabolic tests (GTT and ITT) were performed. HFD-fed mice displayed significantly higher blood glucose levels at all time points during both tests (P<0.05, Fig. 1A and C). As a result, these mice had higher AUC in GTT (44.9%) and lower inverse AUC in ITT (−46.4%) in comparison to LFD-fed mice (P < 0.05, Fig. 1B and D), indicating glucose intolerance and insulin resistance phenotypes. Ten-weeks of HIIT did not attenuate glucose intolerance in the HFD+HIIT group compared to the HFD group (Fig. 1A and B). Regarding ITT, although no statistical significance of inverse AUC in ITT was found between HFD+HIIT and HFD groups, the HFD+HIIT group exhibited a 52% greater AUC in ITT than the HFD group and had no difference when compared to the LFD group (Fig. 1D).

**FIGURE 1.**
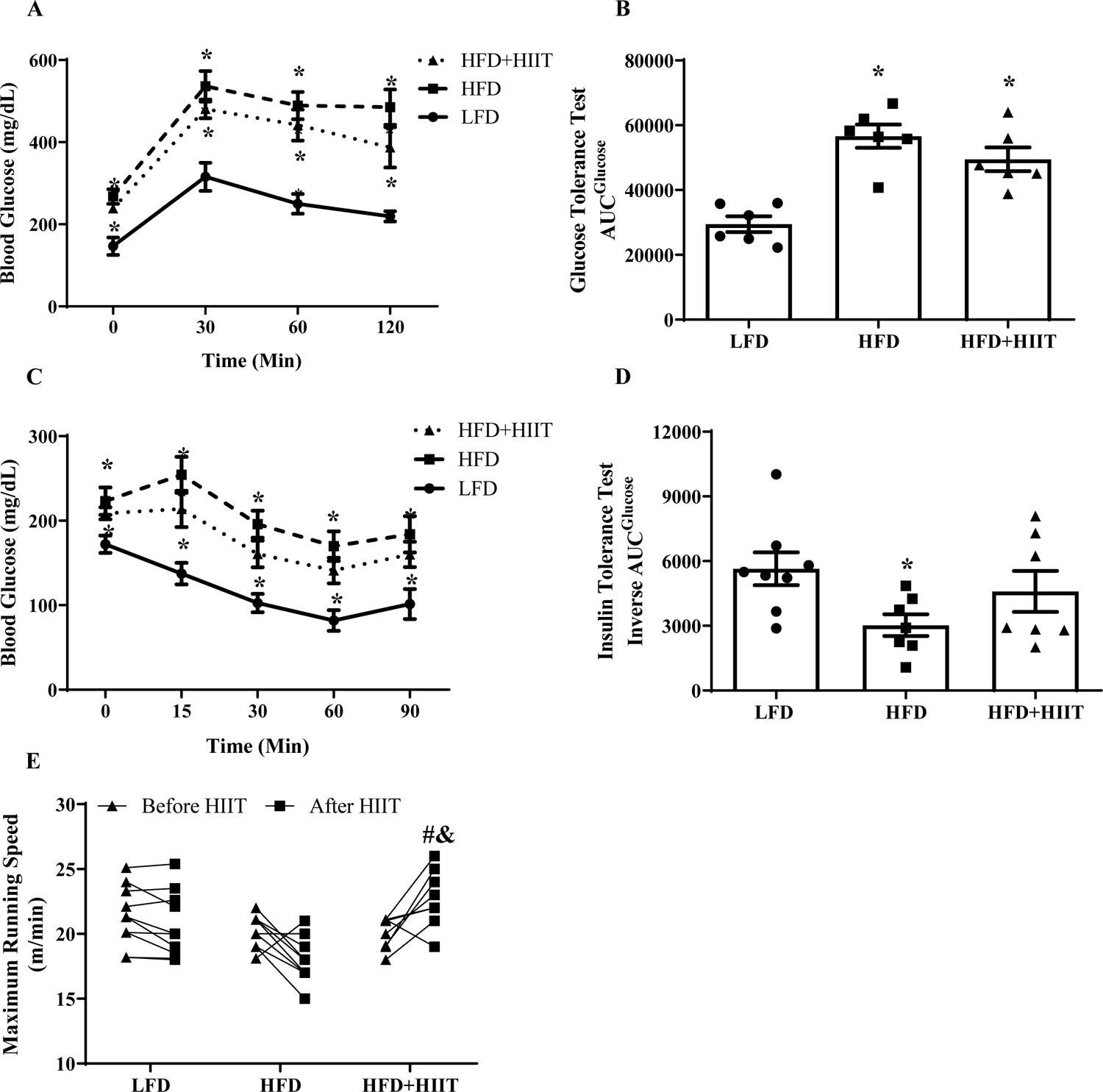
Whole-body glucose homeostasis, insulin sensitivity and exercise capacity in LFD- and HFD-fed mice with or without HIIT. (A) Blood glucose response to GTT. (B) Blood glucose area under the curve (AUC) analysis from GTT. (C) Blood glucose response to ITT. (D) Inverse AUC analysis from ITT. (E) Maximal running speed during Graded Exercise Test (GXT) pre and post HIIT intervention. N = 6-9/group. *P < 0.05 vs LFD, & P < 0.05 vs. pre HIIT intervention. Data are expressed as Mean ± SEM.

Exercise testing revealed that maximal speed increased following 10 weeks of HIIT compared to their pre-training state (11.2%, P < 0.05, Fig. 1E). Further, 10-weeks of HIIT led to significantly higher maximum speeds than LFD and HFD groups (13.0% and 30.7%, respectively. P < 0.05, Fig. 1E).

### Ten Weeks of HIIT improved mitochondrial respiration but did not alter mitochondrial content in skeletal muscle from HFD-induced obese mice

No differences in State 2 (basal pyruvate/glutamate/malate-induced respiration) and State 4 (Olygomycin, an ATP synthase inhibitor) respirations rates were found among groups (Fig. 2A and B). However, State 3 (ADP-stimulated) respiration was significantly lower in the HFD group than LFD group (P < 0.05, Fig. 2A and B). Interestingly, 10 weeks of HIIT markedly enhanced State 3 respiration in skeletal muscle from HFD-fed mice (P < 0.05, Fig. 2A and B). There were no differences in the expression of protein markers of mitochondrial content (OXPHOS, Citrate Synthase, and VDAC) among groups (Fig. 2C-F).

**FIGURE 2.**
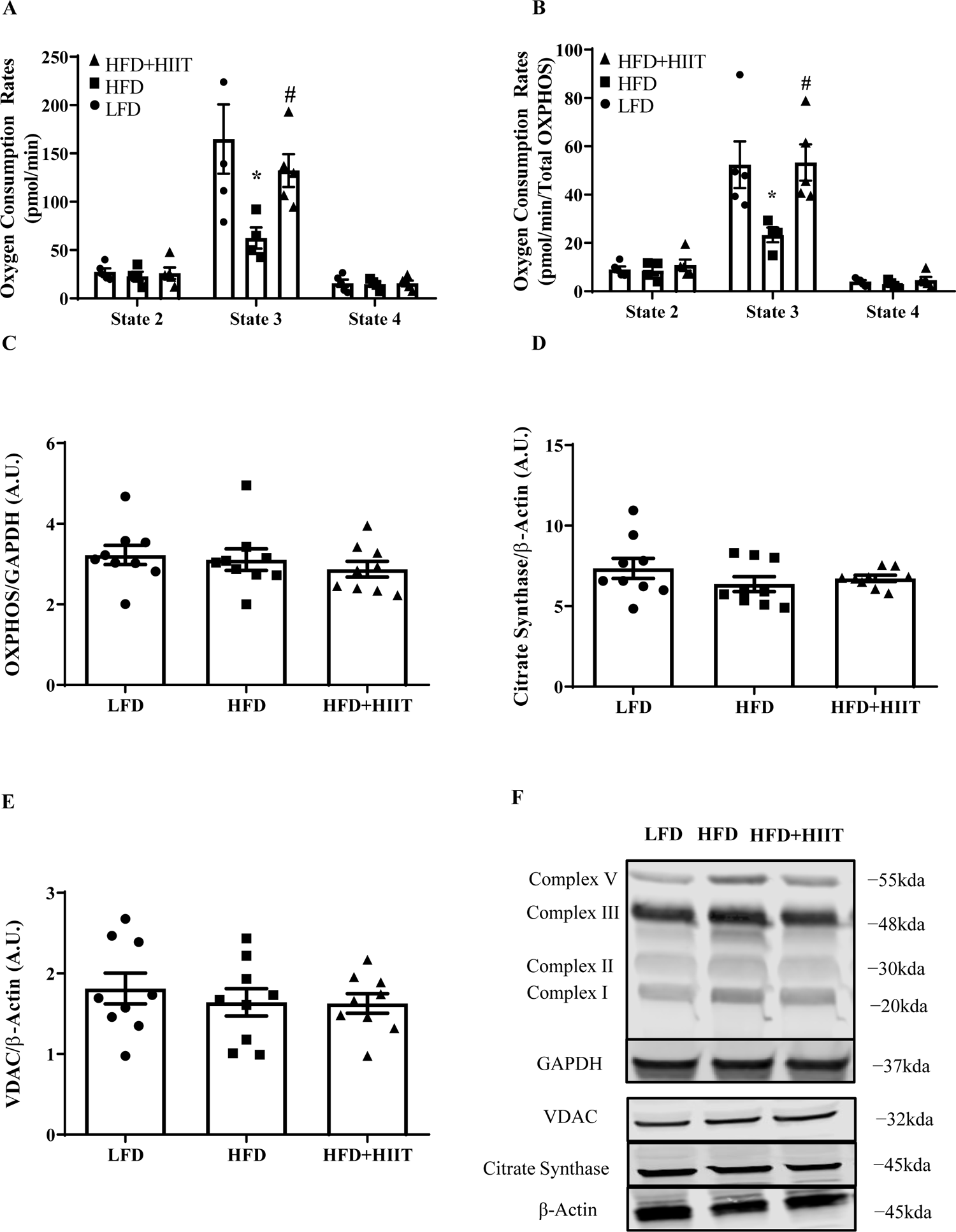
Skeletal muscle mitochondrial respiration and content in LFD- and HFD-fed mice with or without HIIT. (A) Mitochondrial oxygen consumption rates under different respiration states energized by pyruvate+malate. N=4/group. (B) Mitochondrial oxygen consumption rates after normalizing to total oxidative phosphorylation complexes (OXPHOS) content. N=4/group. (C) OXPHOS protein content. (D) Citrate synthase (CS) protein content, (E) mitochondrial voltage-dependent anion channels (VDAC) protein content. (F) Representative immunoblots for C-E. N = 9/group. *P < 0.05 vs LFD, # P < 0.05 vs HFD. Data are expressed as Mean ± SEM.

### Effects of HIIT on protein markers of mitochondrial dynamics in skeletal muscle from HFD-fed obese mice

Skeletal muscle Mfn1 protein content, a regulatory protein of mitochondrial fusion, was significantly lower in both HFD and HFD+HIIT compared to LFD (−55.5% and −41.1%, respectively, P < 0.05, Fig 3A). There were no differences in the contents of other mitochondrial fusion proteins (i.e., Mfn2 or Opa1) among groups (Fig 3B-C).

**FIGURE 3.**
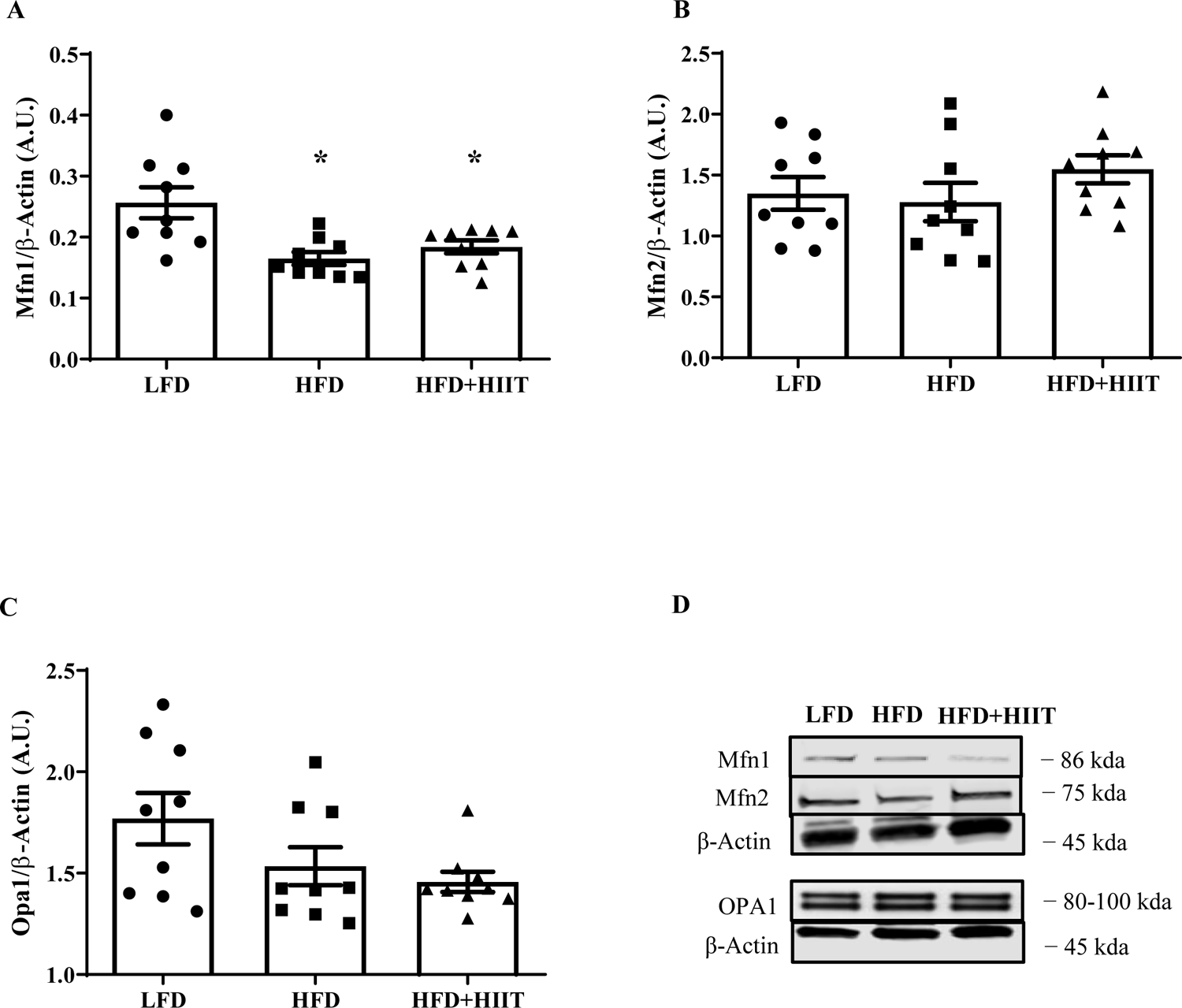
Protein expression of skeletal muscle mitochondrial fusion regulatory proteins in LFD- and HFD-fed mice with or without HIIT. (A) Mfn1 protein content. (B) Mfn2 protein content, (C) Opa1 protein content. (D) Representative immunoblots for proteins in A-C. N = 9/group. *P < 0.05 vs LFD, # P < 0.05 vs HFD. Data are expressed as Mean ± SEM.

With regard to mitochondrial fission, there were no significant differences in skeletal muscle Drp1 protein content among groups (Fig. 4A). However, phosphorylation of Drp1(Ser^616^), a positive regulatory site of Drp1 activity, was higher in the HFD group (40.7%, P < 0.05) compared to LFD group. Interestingly, the increase in phosphorylation of Drp1(Ser^616^) was not exhibited in the HFD-HIIT group (Fig. 4B). Consistently, phosphorylation of Drp1(Ser^637^), a negative regulatory site of Drp1 activity, was significantly lower in skeletal muscles from HFD group when compared to LFD (−87.2%, P < 0.05), but not in skeletal muscles from HFD-fed mice that received 10 weeks of HIIT (Fig. 4C). We further calculated the ratio of phospho-Drp1(Ser^616^) to phospho-Drp1(Ser^637^) as a marker of Drp1 activity. The ratio of phospho-Drp1(Ser^616^) to phospho-Drp1(Ser^637^) was greater in the HFD group in comparison to the LFD group (P < 0.05, Fig. 4D). However, 10 weeks of HIIT reduced this ratio (P < 0.05, Fig. 4D). There was no difference of the ratio of phospho-Drp1(Ser^616^) to phospho-Drp1(Ser^637^) between LFD and HFD+HIIT groups (Fig. 4D). More importantly, correlation analyses revealed a significant positive correlation between AUC of blood glucose in GTT and the ratio of phospho-Drp1(Ser^616^) to phospho-Drp1(Ser^637^) (r = 0.633, P = 0.011, Fig. 2E) and negative correlation between inverse AUC of blood glucose in ITT and the ratio of phospho-Drp1(Ser^616^) to phospho-Drp1(Ser^637^) (r = 0.514, P = 0.024, Fig. 2F).

**FIGURE 4.**
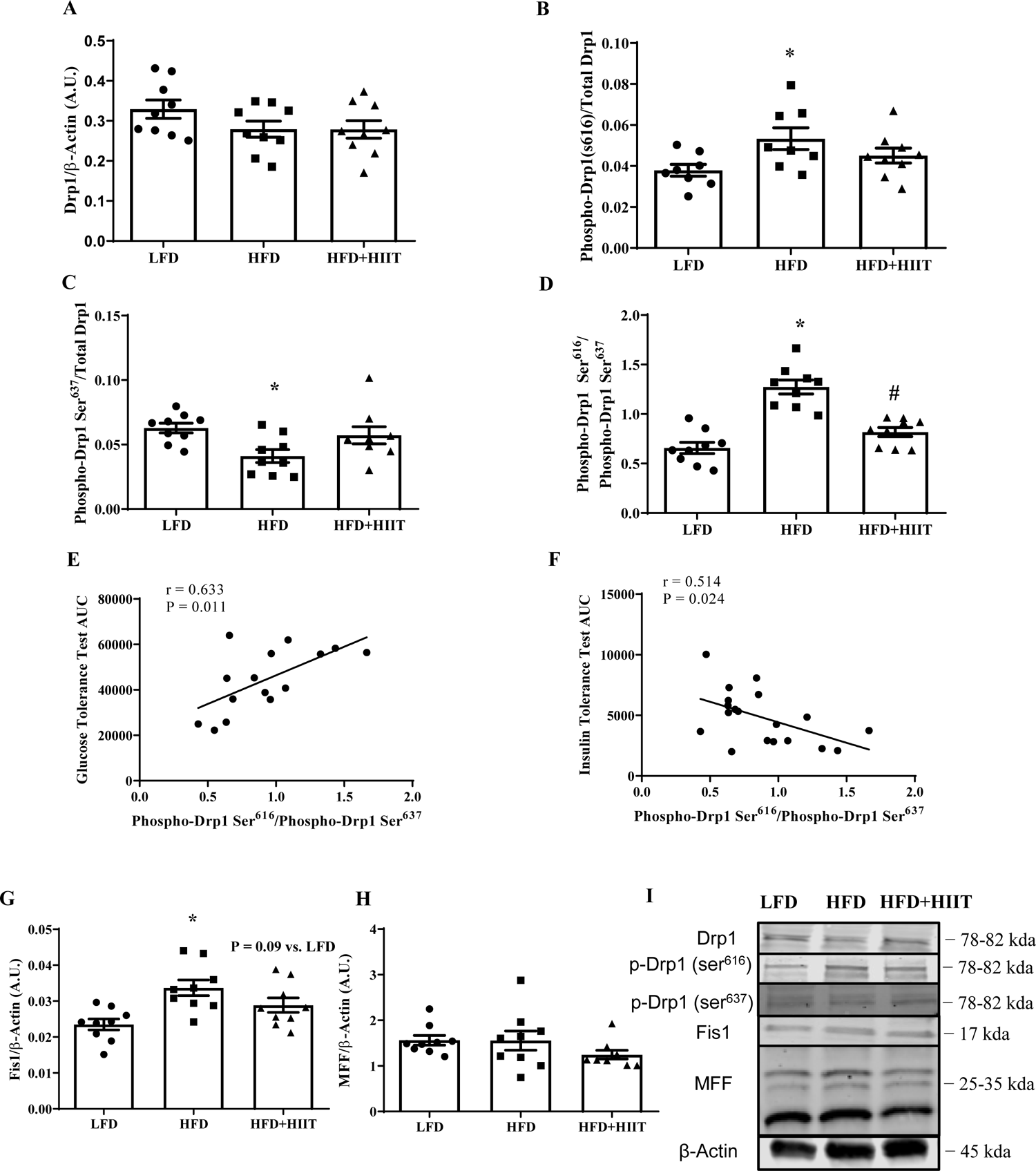
Protein expression of skeletal muscle mitochondrial fission regulatory proteins in LFD- and HFD-fed mice with or without HIIT. (A) Total Drp1 protein content. (B) Ratio of Drp1(Ser^616^) phosphorylation over Total Drp1. (C) Ratio of Drp1 (Ser^637^) over Total Drp1. (D) Ratio of Drp1(Ser^616^) Phosphorylation over Drp1(Ser^637^) phosphorylation. (E) Correlation between the ratio of Drp1(Ser616) Phosphorylation over Drp1(Ser637) phosphorylation and blood glucose area under the curve (AUC) during GTT test. (F) Correlation between the ratio of Drp1(Ser616) Phosphorylation over Drp1(Ser637) phosphorylation and inverse blood glucose area under the curve (AUC) during ITT test. (G) Fis1 protein content. (H) MFF protein content. (I) Representative immunoblots for proteins in A-D and G-H. N = 9/group. *P < 0.05 vs LFD, # P < 0.05 vs HFD. Data are expressed as Mean ± SEM.

We next evaluated the contents of Drp1 adaptor proteins. Specifically, skeletal muscle Fis1 protein content was significantly higher in the HFD group when compared to LFD (30.3%, P < 0.05, Fig. 4G). Interestingly, there is a trend toward a significant reduction of skeletal muscle Fis1 protein content in HFD-fed mice following 10 weeks of HIIT training when compared to their sedentary counterparts (P = 0.09, Fig 4G). There was no difference in Mff protein content among groups.

### Effects of HIIT on protein markers of mitochondrial biogenesis, mitophagy, and autophagy in Skeletal Muscle from HFD-fed obese mice

Although not statistically significant, transcriptional regulator of mitochondrial biogenesis PGC-1α expression was 25% lower in skeletal muscle tissues from the HFD group than LFD group (P = 0.078, Fig. 5A). There was no difference in Parkin, a protein marker of mitophagy, among groups (Fig. 5B). With regard to autophagy, skeletal muscle p62 protein content was significantly lower in HFD group (−35.1%, P < 0.05, Fig. 5C) when compared to LFD group. However, 10 weeks of HIIT increased skeletal muscle p62 protein content in HFD-fed mice (20.8%, P < 0.05, Fig. 5C). Furthermore, LC3B II expression and LC3B II/I were increased in HFD mice compared to LFD mice (34.2% and 15.5% respectively, P < 0.05, Fig. 5E&F), but were significantly attenuated in HFD+HIIT group (−28.6% and −29.9%, P < 0.05, Fig. 5E & F). In addition, the HFD+HIIT group also showed trends toward significance in LC3B II expression and LC3B II/I compared to the LFD group (P = 0.082 and 0.064, respectively, Fig. 5E & F). No differences in LC3B I were found among all three groups (Fig. 5D).

**FIGURE 5.**
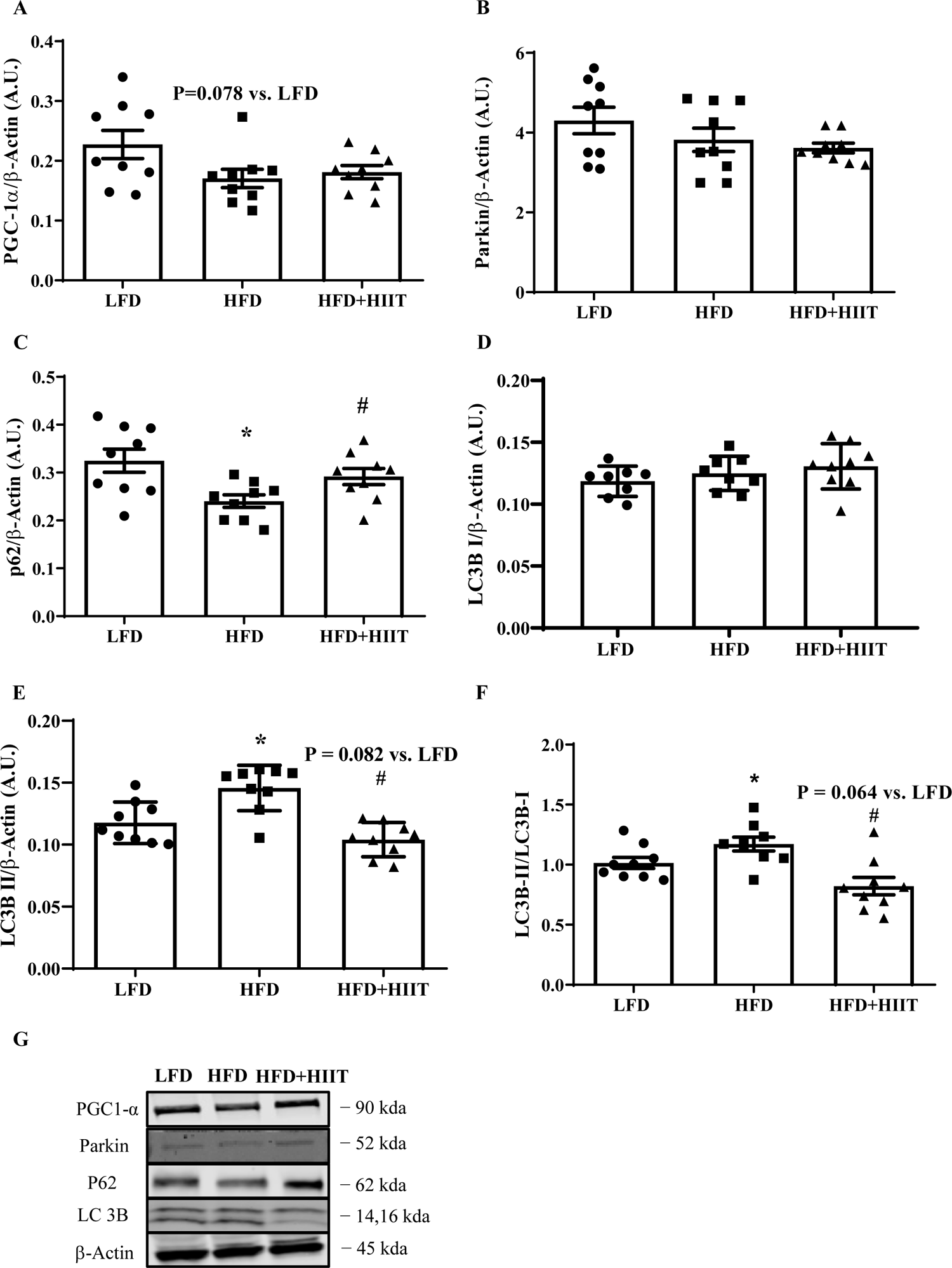
Protein expression of skeletal muscle mitochondrial biogenesis, mitophagy and autophagy regulatory proteins in LFD- and HFD-fed mice with or without HIIT. **(A)** PGC1-α protein content. (B) Parkin protein content. (C) p62 protein content. (D) LC3B Iprotein content. (E) LC3B II protein content. (F) LC3B II:I. (G) Representative immunoblots for proteins in A-D. N = 9/group. *P < 0.05 vs LFD, # P < 0.05 vs HFD. Data are expressed as Mean ± SEM.

## DISCUSSION

The focus of this study was to examine the effects of a HIIT intervention on regulatory proteins governing skeletal muscle mitochondrial quality control processes in diet-induced obese mice. The main finding of the current study was that 10 weeks of HIIT was effective in improving skeletal muscle mitochondrial respiration and the regulatory machinery of mitochondrial quality control, specifically by reducing mitochondrial fission protein Drp1 activity and enhancing autophagy machinery (p62 and LC3B) in diet-induced obese mice. Furthermore, our correlation analyses revealed that alterations in mitochondrial fission protein Drp1 activity was highly associated with responses in glucose and insulin tolerance tests, suggesting whole-body glucose homeostasis and insulin sensitivity may be regulated, at least partly, by the balance of skeletal muscle mitochondrial dynamics.

In the current study, we found that 10 weeks of HIIT partially rebalanced skeletal muscle mitochondrial dynamics towards pro-fusion in diet-induced obese mice, mainly via reducing Drp1-mediated mitochondrial fission. Consistent with previous studies from both humans and animal models (Axelrod et al. 2021; Fealy et al. 2014; Heo et al. 2021; Kugler et al. 2020; Kugler et al. 2021a; Kugler et al. 2021b; Leduc-Gaudet et al. 2018), we found HFD feeding induced excessive activation of Drp1, which was associated with insulin resistance. While few studies have investigated the effects of prolonged (≥ 10 weeks) HIIT on mitochondrial fission proteins in skeletal muscle from obese animals or humans, prior research has consistently reported that continuous endurance-type exercise training effectively reduced mitochondrial fission through the deregulation of fission protein content/activity (e.g., Drp1 and Fis1) (Axelrod et al. 2019; Fealy et al. 2014; Heo et al. 2021). Phosphorylation of Drp1(ser616) promotes mitochondrial fission, whereas phosphorylation of Drp1(ser637) inhibits mitochondrial fission (Ko et al., 2018). The present finding of reduced activity of Drp1 (i.e., downregulation of the ratio of Ser616/Ser637 phosphorylation) agrees with these previous studies (Axelrod et al. 2019; Fealy et al. 2014; Heo et al. 2021), suggesting prolonged exercise training, regardless of exercise intensity, is effective in reducing the excessive mitochondrial fission in skeletal muscle under obese and insulin resistant condition. In contrast, Lifang et al. reported that 8 weeks of HIIT significantly increased Drp1 and Fis1 protein contents, an indication of enhanced mitochondrial fission (Zheng et al. 2020). It should be noted that this study used Streptozotocin-induced type 2 diabetic mice, whereas our study used diet-induced obese, non-diabetic mice. The discrepant results from these two studies suggest HIIT may elicit different adaptations in skeletal muscle mitochondrial fission proteins in different disease models.

No alterations occurred in mitochondrial fusion proteins (i.e., Mfn1, Mfn2, and Opa1) following 10 weeks of HIIT in diet-induced obese mice. To our knowledge, no other studies have assessed mitochondrial fusion proteins in skeletal muscle diet-induced obese mice following prolonged HIIT intervention. MacInnis et al., investigated the effects of short-term (2 weeks) HIIT on skeletal muscle mitochondrial adaptations in lean, healthy humans and found that Mfn2 protein content was markedly increased (MacInnis et al. 2017). The disagreement between our study and this study may be due to the duration of the training since different durations of training resulted in different mitochondrial adaptations (Granata et al. 2018). Alternatively, the discrepancy may also be due to the difference in metabolic health status (i.e., lean healthy vs. obese insulin resistant). Interestingly, while we did not directly compare the effects of HIIT and MICT on mitochondrial dynamics proteins in our study, our findings in mitochondrial fusion proteins are largely different from a previous study utilizing MICT (60%-75% of VO_2_max) as a training modality (Heo et al. 2021). Fully rescued mitochondrial fusion protein contents in skeletal muscle were reported in diet-induced mice following 12 weeks of MICT. Taken together, it suggests that the duration and/or total volume of exercise training in the current study may not be sufficient in mitigating impairment in skeletal muscle mitochondrial fusion. Future studies should be warranted to investigate a longer duration of HIIT (> 10 weeks) on skeletal muscle mitochondrial dynamics under the obese condition.

In the present study, we noted a decrease in p62 protein content and an increase of LC3B II/I ratio in diet-induced obese mice when compared to LFD-fed counterparts. LC3B and p62 are commonly used autophagy markers for monitoring autophagic flux(Jiang and Mizushima 2015). p62 is a classical autophagic cargo adaptor that binds directly to LC3B to recruit ubiquitinated proteins to the autophagosome (Kageyama et al. 2021). A decrease in p62 and an increase in LC3B II/I typically reflects a higher rate of autophagic flux (Bjorkoy et al. 2009). Therefore, our findings suggest that autophagy was elevated in skeletal muscle following 20 weeks of HFD feeding, which are consistent with previous studies with similar length of HFD feeding (Heo et al. 2021; Lee et al. 2021). Such elevation is not a surprise as we speculate that there was a greater need of removing damaged mitochondria due to excessive activation of mitochondrial fission in skeletal muscle from these mice. More importantly, 10 weeks of HIIT virtually reversed the impaired cellular machinery of autophagy in skeletal muscle from the HFD+HIIT group compared to their sedentary counterparts. Our findings in autophagy are partially in conjunction with prior research which elucidated that short-term HIIT lowered protein abundance of p62 and both LC3B II/I (Batterson et al. 2023), but further extended these findings by demonstrating more robust remodeling in autophagy machinery with long-term HIIT under the condition of obesity. Altogether, these findings further corroborate the notion that different durations of HIIT may promote distinct adaptations in mitochondrial quality control and overall skeletal muscle tissue remodeling. Interestingly, we did not see any alteration in mitophagy markers (e.g., Parkin). Similarly, Batterson et al. also reported no change of mitophagy protein markers in skeletal muscle from lean, healthy humans following HIIT, suggesting HIIT training up to 10 weeks was ineffective in improving mitophagy machinery and it may need a longer duration to elicit such improvement. It should be noted that there are two methodological limitations with regard to mitophagy: 1) Mitophagy markers were only evaluated in whole muscle tissue, but not in mitochondrial fraction; 2) We only evaluated protein markers of autophagy, but did not evaluate autophagic flux per se. It is imperative for future research to incorporate an autophagy/mitophagy flux assay (e.g., Chloroquine treatment) in investigating skeletal muscle autophagy/mitophagy flux in response to HIIT.

Despite these positive adaptations in the mitochondrial quality control process in skeletal muscle and weight loss, there was only a modest improvement in whole-body metabolic health in diet-induced obese mice following 10 weeks of HIIT. These findings agree with two previous investigations of a similar study duration (e.g., 10-12 weeks) in which HIIT was sufficient in weight loss and improving exercise capacity in obesity, but it may not be sufficient in mitigating glucose intolerance and insulin resistance in diet-induced obesity (Martinez-Huenchullan et al. 2019; Wilson et al. 2018). In contrast, Martinez-Huenchullan et al. reported that 10 weeks of HIIT significantly improved blood glucose AUC in the ITT test in diet-induced obese mice (Martinez-Huenchullan et al. 2019). The findings of Martinez-Huenchullan et al. agree with a large body of evidence identifying improved insulin sensitivity in diet-induced obese mice following prolonged HIIT (Chavanelle et al. 2017; Denou et al. 2016; Wang et al. 2017; Zheng et al. 2020). In these studies, each HIIT session lasted at least 40 minutes, ranging from 8 to 12 bouts of high-intensity exercise (e.g., ≥ 90% maximal speed) (Chavanelle et al. 2017; Denou et al. 2016; Wang et al. 2017; Zheng et al. 2020), whereas our study only administered 30 minutes (6 bouts) per session. We speculate that 30 minutes per session of HIIT may not be a sufficient volume to strongly enhance whole-body insulin sensitivity and glucose homeostasis under obese insulin-resistant conditions. There was one other study that used the same HIIT regimen but 40 minutes per session in diet-induced obese mice and failed to see any significant improvement in whole-body insulin sensitivity (Martinez-Huenchullan et al. 2018). Taken together, although HIIT has been chosen by many people as one of the most popular training modalities due to its time-efficient nature, it may still need to reach sufficient training volume in order to exert benefits in improving whole-body insulin sensitivity and glucose metabolism under the setting of obesity. Future studies should be warranted to compare different training volumes to further optimize the HIIT training regimen to elicit optimal metabolic improvements.

The present study has several limitations. First, we only used male mice in this study. HIIT has been shown to induce sexually dimorphic adaptations (Scalzo et al. 2014; Schmitz et al. 2020). There is a strong need for future research that directly compares HIIT-induced mitochondrial adaptations in male and female individuals. Second, given the need of large yield of isolated mitochondria for mitochondrial respiration assay, we were not able to study the regulatory proteins involved in mitochondrial quality control in the same muscle (i.e., quadriceps). It is unclear whether the regulatory proteins of mitochondrial quality control in different muscles have similar responses to HFD or HIIT. Previous studies have reported distinct alterations in skeletal muscle mitochondria between glycolytic and oxidative muscles in response to HFD and HIIT (Leduc-Gaudet et al. 2018; Ramos-Filho et al. 2015). Therefore, future investigations are needed to extend our findings to other muscles (e.g., quadriceps, soleus, tibialis anterior muscle) in order to determine if similar adaptations occur across all types of muscles following HIIT.

In conclusion, our data demonstrated that high-fat diet feeding induced insulin resistance was associated with impaired mitochondrial respiration and regulatory machinery of mitochondrial quality control processes and these impairments in skeletal muscle were effectively attenuated by 10 weeks of HIIT, specifically by reduced activity of mitochondrial fission protein Drp1 and enhanced autophagy machinery (p62 and LC3B). Despite these positive adaptations in mitochondria following 10 weeks of HIIT, our study failed to observe significant improvement in whole-body insulin sensitivity, suggesting rebalanced skeletal muscle mitochondrial quality control processes may precede significant whole-body metabolic improvements during prolonged HIIT intervention.

## AUTHOR CONTRIBUTION STATEMENT

K.Z. and J.B.T. did the conception and design of the research; J.B.T., B.A.K. and H.S. performed the experiments; J.B.T., B.A.K. and K.Z. analyzed the data, interpreted the results of the experiments, prepared the Figures and drafted the manuscript; J.B.T., B.A.K, H.Y., T.Y. and K.Z. edited and revised the manuscript; all authors approved the final version of the manuscript.

## DATA AVAILABILITY STATEMENT

Data generated or analyzed during this study are provided in full within the published article and available upon request.

